# IMPAIRED OLFACTORY NETWORK FUNCTIONAL CONNECTIVITY IN PARKINSON’S DISEASE: A NOVEL MARKER FOR DISEASE PROGRESSION

**DOI:** 10.1101/2021.01.21.427682

**Authors:** Prasanna Karunanayaka, Jiaming Lu, Mechelle M. Lewis, Rommy Elyan, Qing X. Yang, Paul J. Eslinger, Xuemei Huang

**Author notes:** **Address correspondence to**: Prasanna R. Karunanayaka, Ph.D., Department of Radiology (Center for NMR Research) Penn State College of Medicine, The M. S. Hershey Medical Center, 500 University Dr., Hershey, PA 17033, USA.

## Abstract

**Objective:** Determine the neural basis of olfactory impairment in akinetic-rigid (PD_AR_) and tremor predominant (PD_T_) Parkinson’s disease subtypes.

**Methods:** We combined resting-state fMRI (rs-fMRI) with seed based functional connectivity (FC) in order to delineate the olfactory network’s functional connectivity (ON FC) between PD_AR_ and PD_T_ patients. We then contrasted their ON FC patterns with cognitively normal (CN) subjects. All three groups were closely matched in age, demographic variables, and adjusted for relative cognitive performance. Olfactory function was measured using the University of Pennsylvania Smell Identification Test (UPSIT).

**Results:** UPSIT scores were lower in akinetic-rigid vs tremor subtypes; ON FC values were lower in PD_AR_ compared to PD_T_ and CN, and followed the trend observed in UPSIT scores. UPSIT scores and ON FC values were significantly correlated, reflecting the effects of PD pathologies.

**Conclusions:** The results show that olfactory function differs between PD_AR_ and PD_T_ suggesting a correlation between PD-related motor symptoms and olfactory deficits. ON FC differences accounts for the impaired olfactory functions observed between PD_AR_ and PD_T_. PD_AR_ is known to have worse clinical outcomes and faster cognitive decline compared to PD_T_; therefore, PD-related olfactory dysfunction may serve as a novel metric for enhancing PD prognosis.

## INTRODUCTION

Parkinson’s disease (PD) is traditionally characterized by tremors, rigidity, and bradykinesia [1, 2]. These motor symptoms have been used to identify akinetic-rigid (PD_AR_) and tremor predominant (PD_T_) subtypes [3–5]. PD_AR_ has worse clinical outcomes and faster cognitive decline compared to PD_T_ [6–11]. PD motor symptoms; however, can be preceded by a set of heterogeneous non-motor symptoms, by several years [12]. Impaired olfaction is one such symptom, being a known harbinger of cognitive decline and neurodegeneration in early PD [13–21].

PD motor symptoms can be traced to a loss of dopaminergic cells in the nigrostriatal pathway [22, 23]. Olfactory impairment is a common premotor symptom in *de novo* patients and is associated with extranigral pathology [24–27]. The existing literature remains unclear about the relationships between olfactory and motor symptoms, even though both deficits are predictive of PD dementia [12, 28].

This paper explores the possibility that the behavioral and neural substrates of olfactory function are differentially affected in PD_AR_ and PD_T_ [29]. The University of Pennsylvania Smell Identification Test (UPSIT) was used to measure olfactory function, and resting state fMRI functional connectivity (FC) was used to investigate its neural basis. Considering that there are worse clinical outcomes for PD_AR_ compared to PD_T_, we hypothesized that the olfactory network’s functional connectivity (ON FC) would differ between each subtype. We predicted that the UPSΓT scores and ON FC estimates would be correlated, reflecting differential PD pathophysiology. The results of this study suggests that PD-related olfactory impairment may serve as a promising clinical biomarker—indicating disease severity.

## METHODS

### Study subjects

17 PD_AR_, 15 PD_T_, and 24 cognitively normal (CN) subjects were recruited for the study; subjects were balanced according to the values and measurements given in **Table 1**. PD_AR_ and PD_T_ subjects were additionally matched for disease severity. Both, the PD cohort and the CN subjects (spouses and relatives of PD subjects) were recruited from a tertiary care, movement disorder clinic, and were part of an ongoing PD study at the Hershey medical center. Previously, in Karunanayaka et al. (2016), resting state fMRI data from this particular cohort was used to delineate differences in the default mode network (DMN) between PD_AR_ and PD_T_ [30]. The research question(s) addressed in this manuscript do not overlap and any discrepancies have been highlighted in the *Discussion* section. Protocols of the study strictly followed the principles of the Declaration of Helsinki [31] and were also approved by the Institutional Review Board (IRB) of the Penn State Hershey Medical Center. As stipulated in the IRB approval, informed, written consent was obtained from each subject before taking part in this study.

**Table 1.** Demographic information of the study cohort (mean ± standard deviation). P-values were derived from Fisher’s Exact test for Gender. All other comparisons: Age, Education, Mini-Mental Status Examination (MMSE), and Hamilton Depression Rating Scale (HMD Scale), were performed using one-way analysis of variance. Statistical significance was evaluated using p<0.05. [UPDRS = Unified Parkinson’s Disease Rating Scale III; T/AR = mean tremor/mean akinetic-rigidity score; LEDD = Levodopa Equivalent Daily Dosage]

|  | **CN**<br>**(n =24)** | **PD<sub>AR</sub>**<br>**(n =17)** | **PD<sub>T</sub>**<br>**(n =15)** | **NC vs.**<br>**PD<sub>AR</sub>** | **NC vs.**<br>**PD<sub>T</sub>** | **PD<sub>AR</sub> vs.**<br>**PD<sub>T</sub>** |
| --- | --- | --- | --- | --- | --- | --- |
| Age (yrs.) | 57.8 ± 7.4 | 59.1 ± 7.4 | 61.7 ± 6.7 | 0.836 | 0.236 | 0.569 |
| Gender (M:F) | 11:13 | 9:8 | 6:9 | 0.876 | 0.423 | 0.724 |
| Education (yrs.) | 15.8 ± 2.4 | 13.9 ± 2.1 | 15.2 ± 3.2 | 0.068 | 0.795 | 0.329 |
| MMSE | 29.6 ± 0.9 | 29.1 ± 1.2 | 29.7 ± 0.5 | 0.257 | 0.959 | 0.221 |
| HMD Scale | 3.7 ± 2.1 | 7.2 ± 3.3 | 6.9 ± 4.0 | 0.0003* | 0.078* | 0.215 |
| Disease duration (yrs.) | NA. | 3.7 ± 4.6 | 3.5 ± 3.5 | NA | NA | 0.91 |
| UPDRS III | 1.2 ± 2.6 | 20.8 ± 12.9 | 17.1 ± 9.5 | <0.0001* | <0.0001* | 0.27 |
| T/AR | NA. | 0.18 ± 0.22 | 2.63 ± 1.77 | NA. | NA. | <0.0001* |
| H&Y stage | NA. | 1.7 ± 0.6 | 1.7 ± 0.6 | NA. | NA. | 0.96 |
| LEDD (mg/day) | NA. | 366 ± 229 | 354 ± 282 | NA. | NA. | 0.88 |

All PD subjects were assessed by a movement disorder specialist (XH) following the criteria outlined in Calne et al. (1992) [32]. Two PD subjects had very mild symptoms and were drug naïve; others were treated with antiparkinsonian medications. PD_AR_ and PD_T_ subjects had no other health issues: neurological disorders, hypothyroidism, vitamin B12 and folate deficiencies, or kidney and liver diseases. Only right-handed PD subjects less than 70 years of age with a Mini-Mental Status Examination (MMSE) Score of 24, who took neither a centrally acting acetylcholinesterase inhibitor nor memantine, were included in this study. CN subjects had no history of neurological or psychiatric disorders, or previous head injuries. CN and PD subjects were pre-screened for possible MRI compatibility complications such as metal implants and claustrophobia.

### PD_AR_ and PD_T_ classification

PD subjects were classified as either PD_AR_ or PD_T_ using the modified ratio developed by Schiess et al., which is based on the UPDRSIII, with a numerical ratio derived from mean tremor and akinetic-rigidity scores [8, 33]. Tremor was assessed using a nine-item scale which included a history of left or right arm tremors (two items); rest tremors of the face, lips, chin and each limb (five items), as well as postural tremors of the right and left upper extremities (two items). The 14-item akinetic-rigidity scale assessed the passive range of motion: rigidity of the neck and at each extremity (five items), rapid opening and closing of the hands (two items), finger tapping (two items), rising from a chair (one item), posture and postural instability (two items), gait (one item), and body bradykinesia (one item). Each item was rated from 0 to 4, with zero representing an absence of symptoms (or normal activity), and 4 indicating the presence of significant symptoms or impairments. The mean of each scale was calculated, and then the ratio (tremor/akinetic rigidity score) was determined. Using this method PD_AR_ subjects will have a ratio ≤ 0.8, whereas PD_T_ subjects will have a ratio ≥ 0.9. In the current study, the average ratio for PD_AR_ subjects was 0.18 ± 0.22 (range 0 – 0.73), and 2.63 ± 1.77 (range 0.9-7.14; **Table 1**) for PD_T_ subjects.

### MRI imaging

MRI Data was acquired on a Siemens 3T MRI system (Magnetom Trio, Siemens Medical, USA) with an 8-channel phased array head coil. Imaging was done while PD subjects were on medication. The rs-fMRI data was acquired with the following parameters: TR / TE / FA = 2000 ms / 30 ms / 90°; FOV = 240 × 240 mm^2^; acquisition matrix = 80 × 80; number of slices = 34; slice thickness = 4 mm, and the number of repetitions = 240. A 3D MPRAGE image was also acquired for volumetric analysis and anatomical overlay. To ensure wakefulness during fMRI scanning, subjects were instructed to relax and keep their eyes open. A high-resolution, T1-weighted, 3D gradient-echo sequence (MPRAGE sequence) was also acquired using the following parameters: TR=2300 ms; TE=2.98 ms; flip angle=90°; FOV=256 x 256 mm^2^; matrix size = 256 x 256; slice thickness = 1 mm (no slice gap); number of slices = 160; voxel size = 1 x 1 x 1 mm^3^.

### Olfactory network (ON)

The coordinates of brain regions in the olfactory network (ON) were selected based on a meta-analysis of fMRI task activation studies [34]. **Figure 1** shows the ON, which includes the piriform cortex (PC) [(−22 0 −14) (22 2 −12)], insula [(−30 18 6), (28 16 8)], and orbitofrontal cortex (OFC) [(− 24 30 −10) (28 34 −12)] [34]. These regions of interest (ROIs) in the ON have been shown to have the highest likelihood of being activated during olfactory stimulation [34]. The time courses from these seed regions are then used to identify correlations among remote brain regions, which constitute the extended intrinsic network. Functional connectivity proximal to seed regions is used to identify the core ON [35–37].

**Figure 1.**
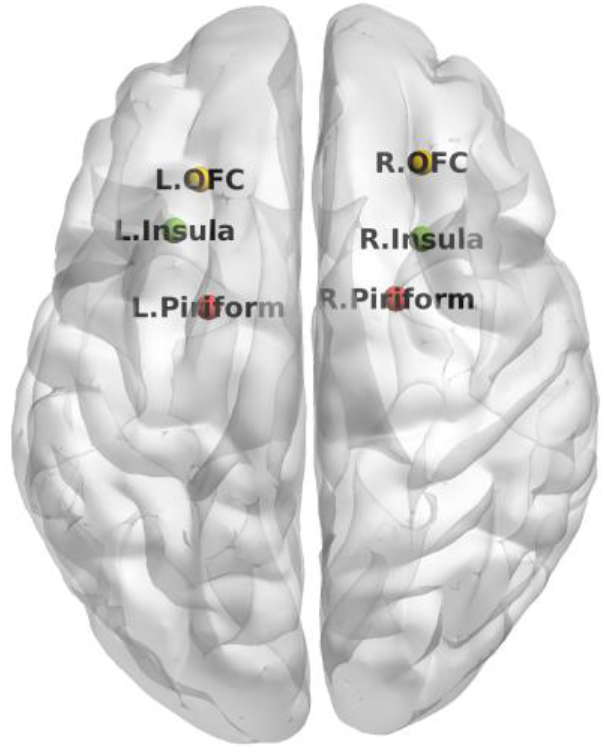
Brain regions in the olfactory network (ON). These regions were selected based on publishedfMRI activation studies. Time courses were extracted in MNI space [(x y z) coordinates] from these six regions as an average time course within a five voxel radius centered on respective coordinates.

### Functional connectivity (FC) analysis

Rs-fMRI data pre-processing and statistical analyses were performed using the DPARSFA (http://rfmri.org/DPARSFA) toolbox. In brief, the methods entailed the following: (1) removal of the first 10 time points; (2) slice time correction; (3) realignment; (4) unified segmentation using 3D-T1 images and spatial normalization using the deformation parameters; (5) time course detrending and spatial smoothing.

The same software was used to estimate motion parameters for all subjects to ensure that they were within 3.0 mm maximum translation for any of the xyz directions, or within 3.0 mm of maximum rotation about the 3 axes. One healthy HC and one PD_T_ had a maximum translation between 2.5 mm and 3.0 mm. We used a seed-based functional connectivity (FC) analysis (detailed in Tobia et al., 2016) to identify differences in ON FC between PD_AR_ and PD_T_. Seed time courses were extracted from pre-processed rs-fMRI data in the Montreal neurological Institute (MNI) space and defined as the average time course within a five-voxel radius centered on the PC, insula, and the OFC coordinates. These time courses were then used to identify correlated brain regions that were remote (i.e., constituents of the extended and intrinsic ON), and proximal to seed regions which constituted the core of the intrinsic ON. Finally, ON FC maps were set to a threshold of p< 0.01 with AlphaSim Correction.

### Voxel-Based Morphometry

Voxel-based morphometry (VBM) was performed using SPM12 (http://www.fil.ion.ucl.ac.uk/spm/). We performed the following steps: (1) inspected T1-weighted anatomical images to ensure no gross anatomical abnormalities; (2) segmented images into grey matter (GM), white matter (WM), and cerebrospinal fluid (CSF), and (3) spatially normalized the segmented images. The image intensity of each voxel was modulated by Jacobian determinants to ensure that regional differences in the total amount of GM volume was conserved; (4) transformed registered images into the Montreal Neurological Institute (MNI) space using the affine spatial normalization. During this normalization step, images were Jacobian-scaled for “modulated” VBM and resampled to 1.5 mm^3^ isotropic voxels; (5) The normalized and modulated GMV images were smoothed with an 8-mm full-width, at half-maximum (FWHM), isotropic Gaussian kernel; (6) A one-way ANOVA was performed to determine volumetric group differences.

### Statistical analyses

Demographic (age and education) and clinical factors (HAM-D, MMSE, LEDD, and UPDRS III) were compared using a simple one-way analysis of variance (ANOVA). The sex ratio between groups was compared using Fisher’s Exact Test. Cognitive test scores were converted to standardized z-scores. One-way analysis of covariance (ANCOVA) was used to examine group differences in each cognitive domain with adjustments for HAM-D scores—where the individual cognitive domain was the dependent variable and the group was the independent variable. One-way ANCOVA was also used to test for group differences in individual cognitive tests with HAM-D scores as a covariate. The Tukey-Kramer method was used to correct for multiple group (3 groups) comparisons. Since multiple cognitive tests were compared, results were corrected for multiple comparisons of five test domains with a false discovery rate (FDR) at the 0.05 level. We report raw p values but indicate whether the results were also significant after FDR correction at a p value of 0.05. Finally, an explorative correlation analysis was performed to test the significance of the relationship between ICA z-scores and the neuropsychological test scores in the five cognitive domains described above.

### Multivariate classification of ON FC values

The discriminative information contained, within the functional connectivity of the olfactory network (ON FC) in PD_AR_ and PD_T_ subtypes, was investigated. A principal component analysis of the multivariate ON FC values (across brain regions in the ON) was combined with the nearest neighbor method, as implemented in Mathematica (Wolfram, Champaign, IL, USA). This approach (sometimes referred to as embedding) incorporates a neurobiological, interpretive, and generative model of the ON—with discriminatory classifiers. These multivariate methods significantly outperform conventional correlation-based FC methods. As stated in Brodersen et al. (2011), these methods can provide accurate separation of networks by exploiting discriminative information encoded in ‘hidden’ physiological quantities, such as synaptic connection strength [38].

## RESULTS

### 3.1. Demographic and cognitive comparisons

There were no significant differences in age, MMSE, sex ratio, or education between PD_AR_, PD_T_, and CN (p > 0.068; see **Table 1** and **Table S1**, Sup. Material). PD subjects had significantly higher UPDRS III scores (p < 0.0001) compared to CN. PD_AR_ subjects had significantly higher HAM-D scores than CN. **Table 2** (below) shows the neuropsychological domains tested.

**Table 2:** Neuropsychological test scores and analyses. Neuropsychological test results showing the mean ± standard deviation. All test scores were converted to standard z-scores. Higher z-scores indicate better performance. One-way analysis of covariance (ANCOVA) was conducted with Group as the independent variable and each Cognitive Domain as the dependent variable, with the Hamilton Depression score serving as a covariate. Individual neuropsychological subtests were also tested using ANCOVA, with the Hamilton Depression score used as a covariate. Here, we report raw p-values. None of the test results were significant after FDR (false discovery rate) correction at a p value of 0.05. [CWInt = Color-Word Interference Test; VVT = Visual Verbal Test; DesFlu = Design Fluency; VerbFlu = Verbal Fluency; JoLO = Judgment of Line Orientation; BVMT-R = Brief Visuospatial Memory Test-Revised; HVLT-R = Hopkins Verbal Learning Test-Revised.] *Discrimination index scores are an index of recognition memory load, i.e., on number of Hits – number of False Alarms.

| Test | CN<br>(*n* = 24) | PD<sub>AR</sub><br>(*n* = 17) | PD<sub>T</sub><br>(*n* = 15) | HC vs.<br>PD<sub>AR</sub> | HC vs.<br>PD<sub>T</sub> | PD<sub>AR</sub> vs.<br>PD<sub>T</sub> |
| --- | --- | --- | --- | --- | --- | --- |
| **<u>Executive function</u>** |  |  |  |  |  |  |
| Z-scores | 0.39 ± 0.40 | 0.05 ± 0.59 | 0.09 ± 0.37 | 0.075 | 0.223 | 0.788 |
| CWInt Inhibition | 0.65 ± 0.70 | 0.29 ± 1.04 | 0.53 ± 0.85 | 0.866 | 0.999 | 0.882 |
| Inhibition Errors | 0.01 ± 0.76 | -0.49 ± 1.03 | 0.33 ± 0.58 | 0.409 | 0.360 | 0.032 |
| CWInt Switch | 0.65 ± 1.00 | -0.08 ± 1.34 | 0.62 ± 0.63 | 0.226 | 0.998 | 0.202 |
| Switch Errors | 0.46 ± 0.64 | -0.08 ± 0.91 | 0.47 ± 0.57 | 0.195 | 0.976 | 0.133 |
| VVT_Total | 0.20 ± 0.80 | -0.21 ± 1.22 | -0.33 ± 1.68 | 0.971 | 0.661 | 0.826 |
| VVT_Switch | 0.05 ± 0.87 | -0.36 ± 1.15 | -0.59 ± 1.77 | 0.978 | 0.524 | 0.686 |
| DesFlu Switch | 0.69 ± 0.89 | 0.55 ± 0.70 | 0.22 ± 0.91 | 0.786 | 0.204 | 0.600 |
| DesFlu Total Correct | 0.82 ± 1.18 | 0.12 ± 0.76 | 0.18 ± 0.81 | 0.286 | 0.234 | 1.00 |
| DesFlu Total Design | 1.07 ± 1.30 | 0.59 ± 0.85 | 0.80 ± 1.42 | 0.739 | 0.897 | 0.943 |
| DesFlu Design Accuracy | -0.36 ± 0.92 | -0.75 ± 1.12 | -0.80 ± 1.43 | 0.531 | 0.452 | 0.998 |
| VerbFlu Letter | 0.01 ± 0.85 | -0.43 ± 1.14 | -0.41 ± 0.85 | 0.101 | 0.191 | 0.900 |
| VerbFlu Category | 0.46 ± 0.92 | 0.29 ± 0.90 | 0.08 ± 1.02 | 0.429 | 0.245 | 0.959 |
| **<u>Spatial cognition</u>** |  |  |  |  |  |  |
| Z scores | 0.15 ± 0.71 | -0.31 ± 0.79 | -0.03 ± 0.58 | 0.183 | 0.755 | 0.505 |
| JoLO | 0.30 ± 1.42 | -0.51 ± 1.50 | 0.05 ± 1.17 | 0.252 | 0.742 | 0.632 |
| DRS2 Construction | 0 | -0.12 ± 0.33 | 0 | 0.243 | 0.997 | 0.214 |
| **<u>Learning/Memory</u>** |  |  |  |  |  |  |
| **<u>Spatial Memory</u>** | 0.03 ± 0.84 | -0.61 ± 1.35 | -0.41 ± 1.48 | 0.695 | 0.768 | 0.985 |
| BVMT Learning | 0.00 ± 1.10 | -0.51 ± 1.24 | -0.48 ± 1.40 | 0.828 | 0.693 | 0.981 |
| BVMT Delayed Recall | 0.21 ± 1.24 | -0.27 ± 1.20 | -0.01 ± 1.32 | 0.975 | 0.999 | 0.977 |
| BVMT Discrimination Index* | -0.10 ± 1.29 | -1.05 ± 2.24 | -0.74 ± 2.14 | 0.558 | 0.715 | 0.951 |
| **<u>Verbal Memory</u>** | -0.57 ± 1.15 | -0.55 ± 1.00 | -0.43 ± 0.66 | 0.982 | 0.959 | 0.898 |
| HVLT Total Learning | 0.49 ± 1.15 | -0.64 ± 1.03 | -0.78 ± 0.92 | 0.717 | 0.571 | 0.983 |
| HVLT Delayed Recall | -0.65 ± 1.25 | -0.68 ± 1.09 | -0.42 ± 0.98 | 0.967 | 0.877 | 0.760 |
| HVLT Discrimination Index* | -0.57 ± 1.47 | -0.33 ± 1.31 | -0.09 ± 0.62 | 0.896 | 0.544 | 0.846 |
| **<u>Attention/working Memory</u>** |  |  |  |  |  |  |
| Z-scores | 0.62 ± 0.68 | 0.29 ± 0.50 | 0.41 ± 0.64 | 0.301 | 0.576 | 0.845 |
| Letter Number Sequencing | 0.43 ± 0.75 | 0.33 ± 0.88 | 0.09 ± 0.82 | 0.818 | 0.357 | 0.765 |
| Spatial Span | 0.44 ± 0.90 | 0.22 ± 0.90 | 0.44 ± 1.16 | 0.913 | 0.990 | 0.855 |
| Digit Span | 0.42 ± 0.68 | 0.20 ± 0.71 | 0.29 ± 0.74 | 0.301 | 0.642 | 0.791 |

### UPSIT group differences

The ANCOVA analysis detected group differences in UPSIT scores (F = 54.21, p < 0.001). **Figure 2** shows the results of a post hoc analysis showing group differences in UPSIT scores. CN showed significantly higher UPSIT scores compared to PD subtypes. PD_AR_, however, had significantly lower UPSIT scores than PD_T_.

**Figure 2.**
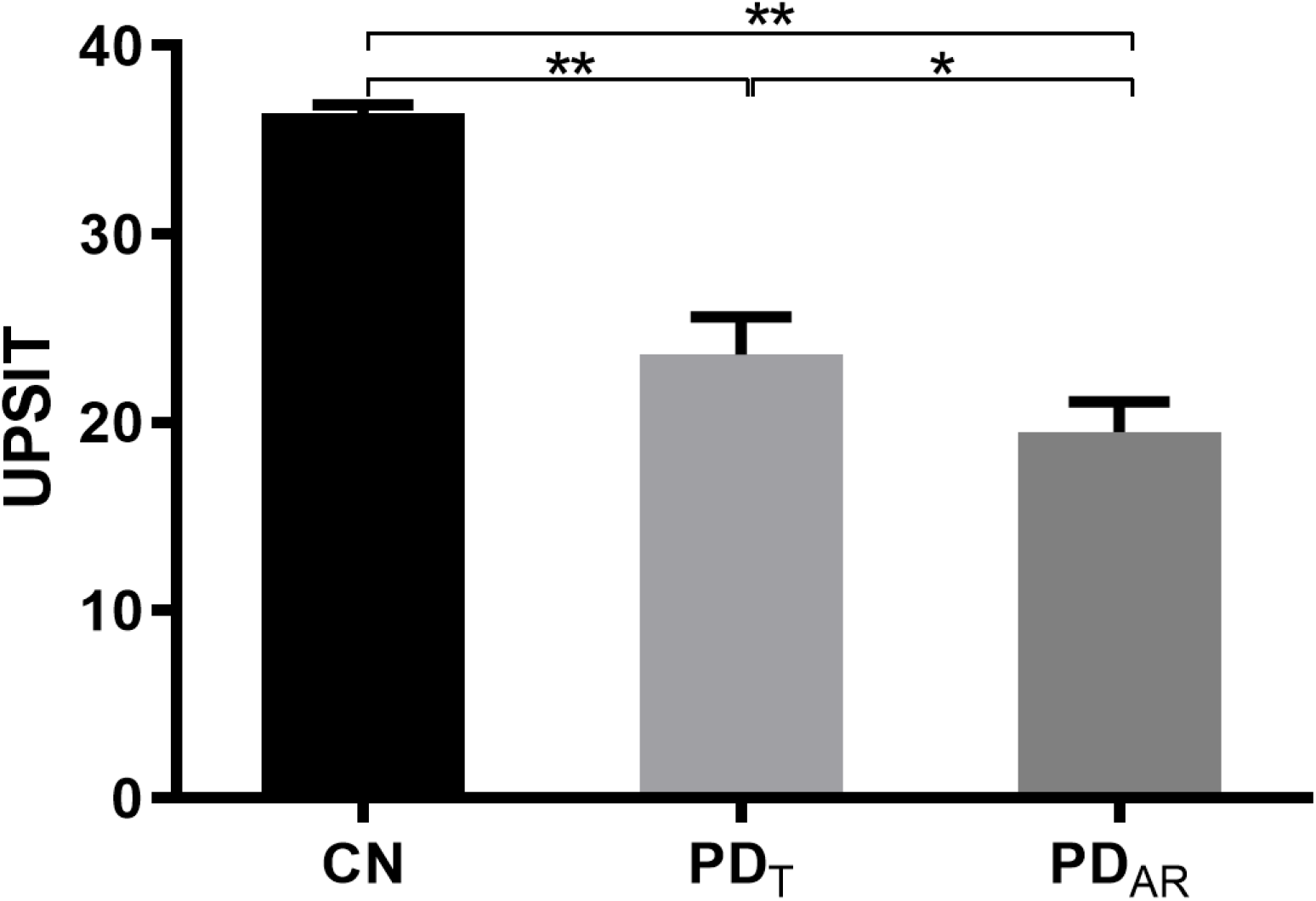
*Group differences in UPSIT scores (mean* ± SD) *between CN, PD_T_ and PD_AR_*. CN: 6.41±2.15; PD_T_: 23.58±6.96; PD_AR_: 19.47±6.65. *,*p <0.05; ** p <0.01*.

### Volume differences

VBM differences between groups were not detected after correcting for multiple comparisons with family wise error (FWE) in SPM12. Differences were detected within the supplementary motor area with a statistical threshold of p < 0.001, uncorrected. A detailed analysis of GM volumes in respective groups is provided in Karunanayaka et al. (2017). Note that this study may be underpowered to detect significant GM group differences between CN and PD.

### ON FC group differences

In order to investigate the neural basis of UPSIT score differences shown in **Figure 2**, we conducted a whole brain one-way ANOVA analysis of ON FC values for each group. The POC, OFC, insula, cerebellum, Hippocampus, and posterior cingulate cortex (PCC) showed ON FC group differences (**Figure 3**).

**Figure 3.**
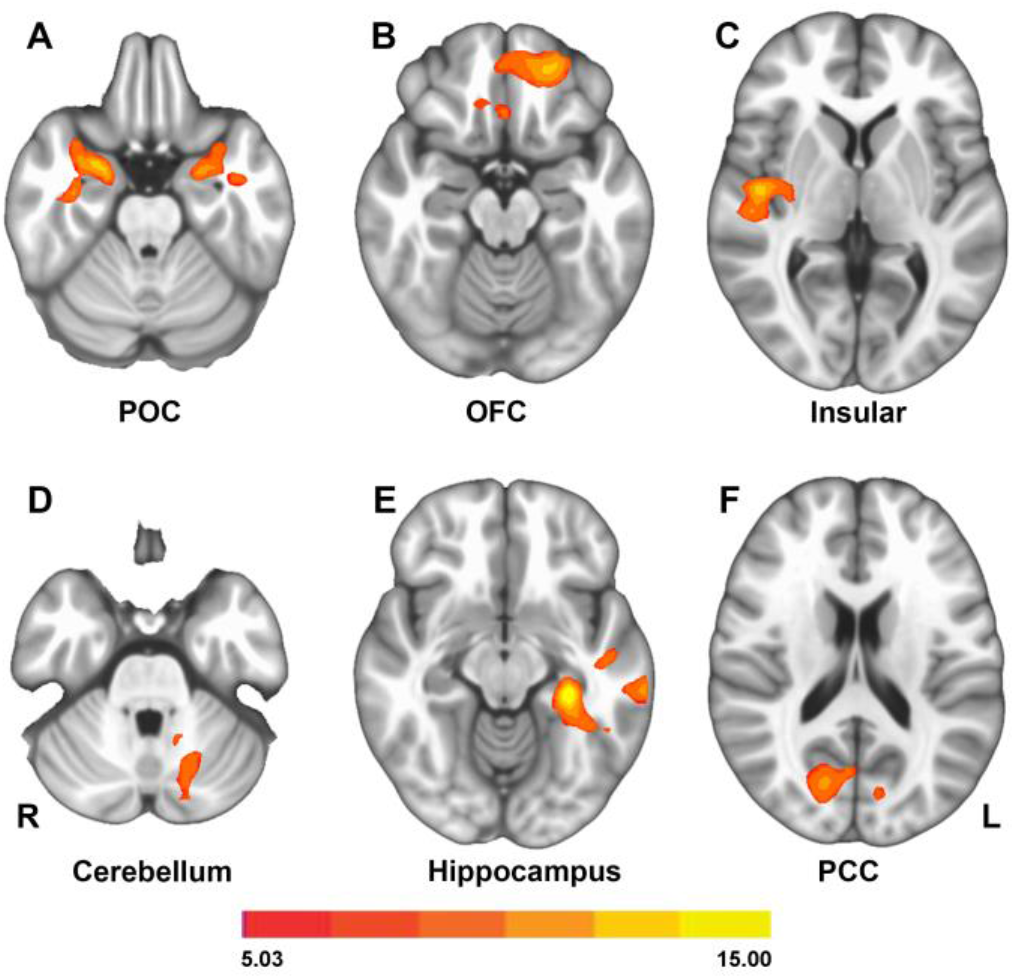
Group differences in Olfactory network functional connectivity (ON FC) between CN (cognitively normal), PD_AR_, and PD_T_groups.p <0.01, AlphaSim Corrected.

We extracted ON FC values within the six brain regions shown in **Figure 3** and investigated their group differences (**Figures 4** and **5**). In the core ON (**Figure 4**), POC connectivity to the ON differs significantly between the CN and both PD subtypes. In contrast, both OFC and insula connectivity to the ON is significantly different between PD_AR_ and CN, as well as PD_AR_ and PD_T_. These results support a differential ON FC patterns between PD_T_ and PD_AR_ within the core ON.

**Figure 4.**
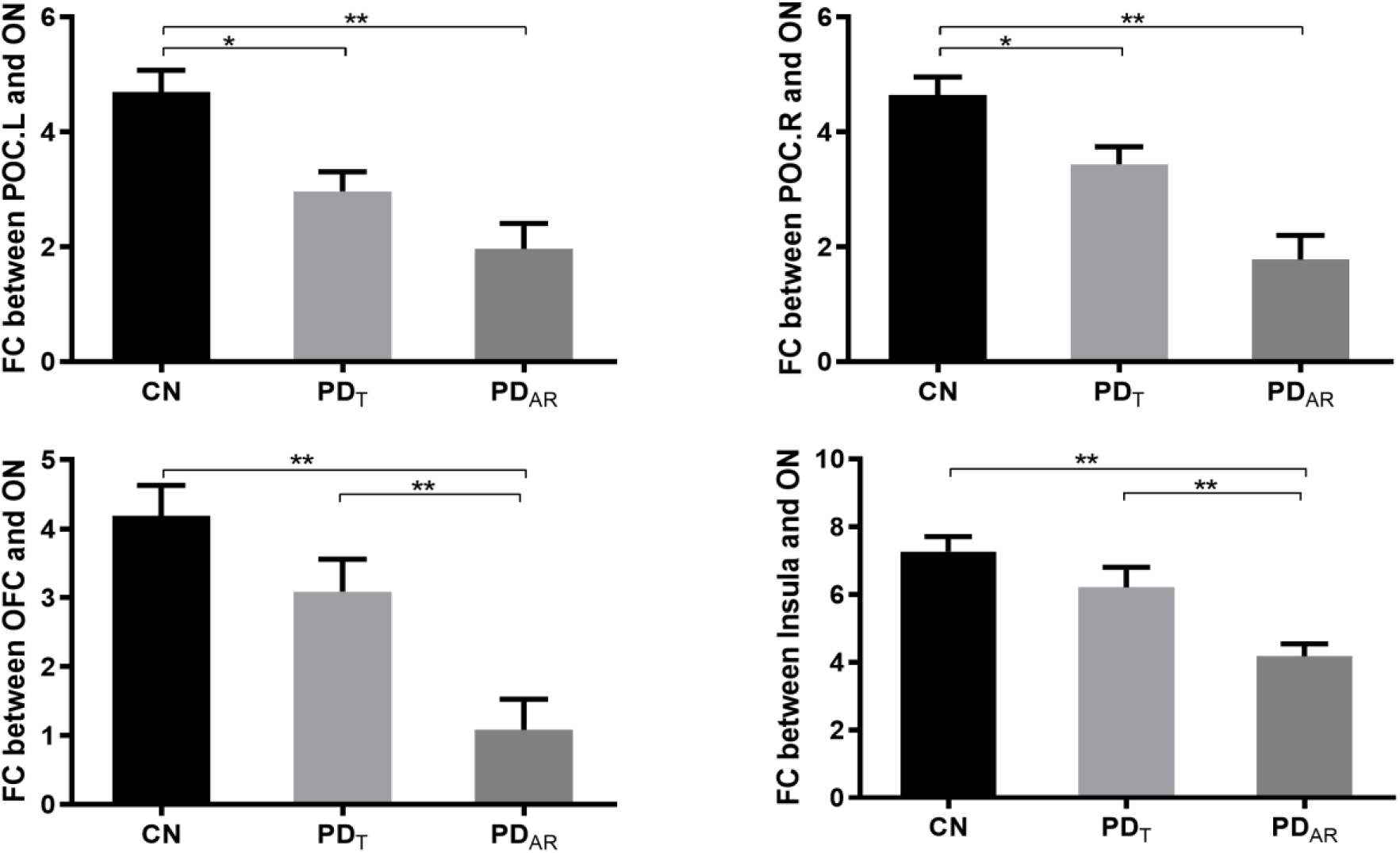
ON FC group differences within the core olfactory network. The ON FC to the OFC *and* insula significantly differs between PD_AR_ and PD_T_. *, p < 0.05; **, p < 0.01.

**Figure 5.**
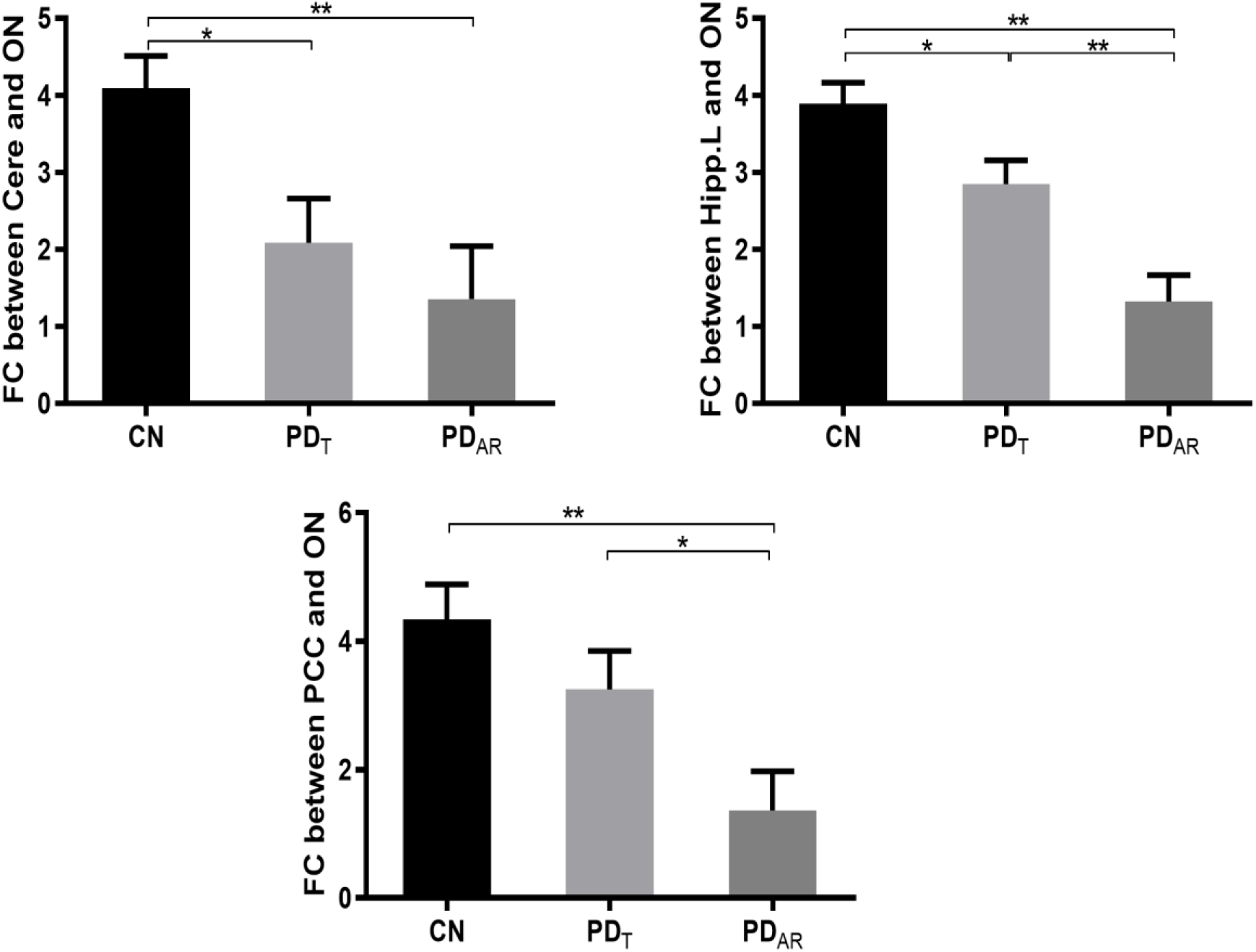
ON FC differences within the extended olfactory network. The FC of the left hippocampus *and* the posterior cingulate cortex (PCC) significantly differs between PD_AR_ and PD_T_. *, p < 0.05, **: p < 0.01. Cere, Cerebellum.

A similar pattern of differential ON FC is observed within the extended ON. Here, ON FC between the left hippocampus and the PCC significantly differs between PD_T_ *and* PD_AR_. Note that the left hippocampus FC to the ON differs among all groups.

### Correlations between ON FC and UPSIT scores

Average ON FC values within the six brain regions shown in **Figure 2** were not correlated with UPSIT scores in respective groups. However, as shown below, ON FC is positively correlated with UPSIT scores in the combined group (**Figure 6**). This observation likely reflects differences between CN and PD, including differences in disease severity between PD_AR_ and PD_T_.

**Figure 6.**
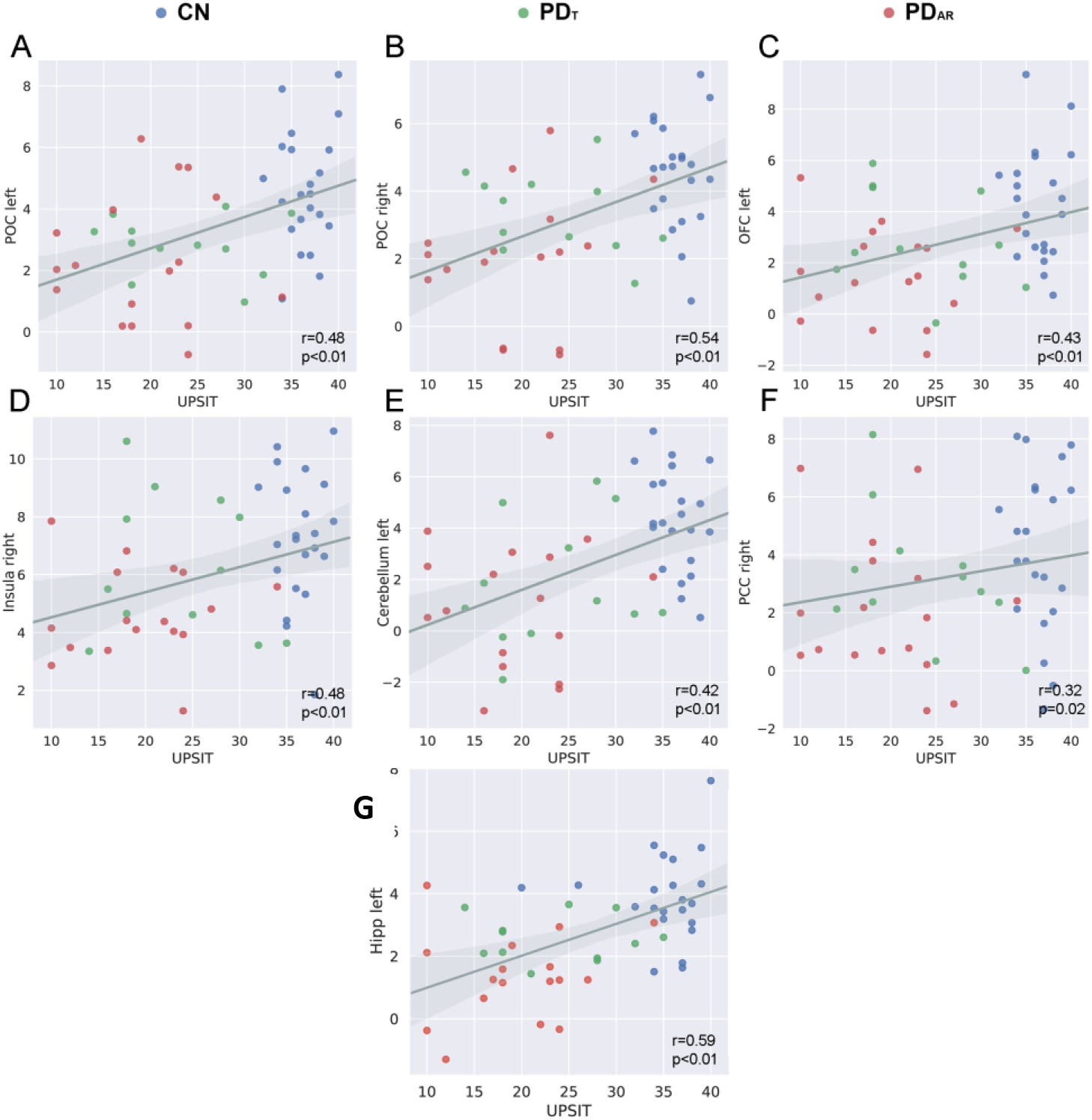
The correlations between UPSIT scores and the FC of each brain region with the ON. Since PD_AR_ shows poor prognosis, these results suggest that ON FC may have the capacity to serve as an indicator of disease severity.

### Multivariate classification of ON FC

The nearest neighbor classification analysis (based on PCA) of multivariate ON FC data is shown in **Figure 7**. The classification accuracy for PD_T_ was 67% and 83% for PD_AR_, suggesting that ON FC contains above chance (i.e., 50%) discriminatory information for separating PD_T_ from PD_AR_.

**Figure 7.**
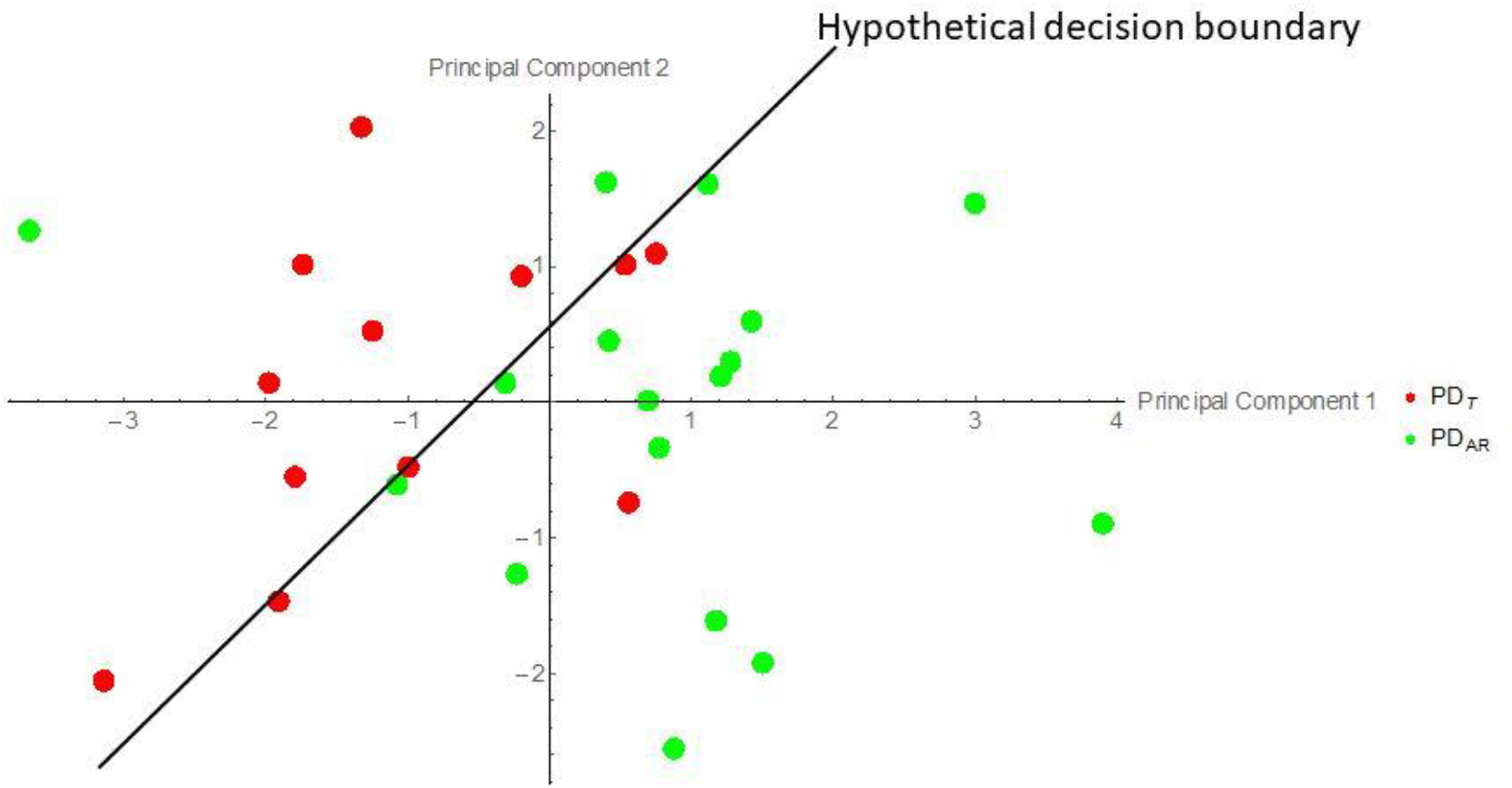
Multivariate ON FC data projected onto the first two principal components. In this reduced plane, a decision boundary can be identified, separating PD_AR_ and PD_T_ at above chance levels.

## DISCUSSION

High prevalence, early presence, persistence throughout disease progression, and ease of olfactory testing, have all increased interest in testing whether olfactory dysfunction can serve as an effective biomarker for the early diagnosis, differential diagnosis, and prognosis of clinical outcomes associated with neurodegenerative diseases. The results of this study strongly support a type-dependent ON FC impairment in PD. It has been suggested that differences in PD-related motor symptoms are indicative of differences in neural networks, which are preferentially impacted by the disease [8, 29]. These PD differences also appear to be related to the clinical progression of PD, such as the risk of cognitive decline leading to dementia [29]. Following this PD pathophysiology, differential olfactory impairments were revealed between PD_T_ and PD_AR_ subjects who were in the early stages of a PD cascade. As hypothesized, PD_AR_ subjects (known for rapid motor and cognitive decline) exhibited lower UPSIT scores and ON FC when compared to cognitively comparable PD_T_ subjects [9]. This type-dependent ON FC strengthens existing evidence, and highlights the deficits in the central olfactory system in PD, as well as the differential neuropathologies between PD_T_ and PD_AR_ [8, 39].

### ON FC distribution

We observed a positive correlation between ON FC and UPSIT scores in the combined group. This shows that ON FC is linked to behavioral olfactory measurements in a meaningful way. Although the within group correlations between the ON FC and olfactory performances (UPSIT scores) were not quantified, the combined group correlations (**Figure 6**) clearly demonstrated type-dependent clustering behavior. These results may help establish specific relationships between ON FC and pathological changes in the POC/hippocampus, as well as ON FC and PD-related behavioral measurements. Similar relationships have previously been hypothesized for AD pathologies using ON FC in the olfactory structures [37]. Our results provide in vivo evidence for olfactory network (ON) involvement—beyond the olfactory bulb and tract—in PD pathology. Altered olfaction in PD seemingly owes its origins to changes in central olfactory processing [40].

Olfactory deficits in early PD are commonplace, and currently serve as crucial diagnostic tools. This deficit manifests as a prodromal symptom, and frequently precedes motor and cognitive symptoms. For example, the reduced intrinsic integrity of the substantia nigra in subjects with unexplained smell loss, has been shown to confirm the PD “at-risk” status [41]. Similarly, being normosmic with normal cognition at time of diagnosis has been shown to be a reliable predictor of stable cognitive functions for up to 10 years [42]. Numerous studies have shown that PD-related olfactory impairments can be used for the differentiation between PD and other movement disorders [43, 44].

Our results raises the possibility that preferential involvement of the ON in PD_T_ and PD_AR_ can produce measurable differences in ON FC *before* the onset of cognitive impairments [30]. In other words, ON FC could serve as a useful prognostic marker when considering the risk of developing cognitive decline [e.g., mild cognitive impairment (MCI)] in PD. The results suggest that ON FC may serve as a useful marker, reflecting disease severity. These findings point to a novel brain-behavior relationship, highlighting a close correspondence between motor and olfactory impairments in PD.

Motor symptoms in PD can be traced back to a loss of dopaminergic cells in the nigrostriatal pathway [45]. On the other hand, PD-related olfactory deficits do not respond to dopaminergic medications and are thought to be caused by extranigral pathologies [46]. Our results suggest that both olfactory and motor impairments of PD share a common, underlying neural substrate; this substrate, if impaired, would increase one’s chances of developing PD dementia.

### Network perspective of olfactory function in PD

It has been suggested that localized functions of brain areas are specified primarily by their distributed global connectivity [47, 48]. Our study extends this idea to olfaction by demonstrating that the ON FC is capable of providing a mechanistic explanation of PD-related olfactory impairment. It should be noted that identifying the neural mechanisms by which PD pathology disrupts brain functions is of critical importance for the development of early diagnostic tools and prophylactics aimed at inhibiting disease progression.

Pathological protein aggregates also affects olfactory brain regions before other regions, making the ON particularly vulnerable to neurodegeneration [43, 49, 50]. Indeed, pathology in the olfactory bulb (OB), which has bidirectional projections to the POC, is common in early stage PD [51]. Note that the OB cannot easily be imaged using fMRI at 3T and, as a result, is not included in the current study. Nevertheless, it has been suggested that the ubiquitous, though varying, olfactory impairments in neurodegeneration may be caused by a differential disruption to a common primordial neurological substrate [9].

### Resting state (RS) fMRI

RS brain networks are intrinsically organized and modulated during task performance [47, 52]. A strong correspondence between RS functional connectivity (RSFC) and task-evoked FC has also been established, which includes the olfactory system. Similarly, a close correspondence has been established between RS brain networks and the brain’s structural connectivity [53–55]. The implemented methodologies in this paper have enabled us to investigate whether the ON FC becomes stronger or weaker within the core and extended ON, and in PD_T_ and PD_AR_. Within the core ON, no significant differences in ON FC to the POC was observed. In contrast, ON FC to the OFC and insula were significantly impaired in PD_AR_. Furthermore, ON FC to the hippocampus and PCC (within the extended network) were significantly lower in PD_AR_. Additionally, the pattern of ON FC to the hippocampus followed the same trends as the UPSIT scores seen in **Figure** 2. These findings corroborate the study by Westermann et al. (2008), which suggested that reduced hippocampal activity had an effect on olfactory sensitivity [56]. Our data shows extensive and complex ON FC changes in PD pathology. Our study suggests that resting state FC changes in the ON, which is a measure of the central nervous system, reflects impairments in olfaction, in PD patients [57].

### Gray matter comparison

In PD, atrophy in the ON and its relation with olfactory impairment remains unclear [58, 59]. Some studies reported cortical gray matter (GM) losses [e.g., linked right piriform cortex (PC) atrophy to olfactory deficit] but others failed to report associations between GM losses and olfactory impairment [60–62]. The present study did examine cortical atrophy in the ON and explored its relationship with olfactory function in a relatively large, early-stage sample of PD patients. The lack of statistically significant GM volume loss, within the ON, may suggest that significant regional brain atrophy might be lower in the absence of comorbid cognitive impairment between PD subtypes. Our results suggest that functional changes in the ON may be more sensitive than GM volume changes. Since PD-related olfactory deficits are predictive of dementia, ON FC may be a useful marker in detecting and predicting cognitive decline risks in PD [63].

### Multivariate ON FC

Our results showed sufficient discriminatory information about are embedded in the multivariate ON FC measurement (1^st^ two principal components of ON FC). For example, the PCA captured sufficient ON FC variation across brain regions, distinguishing PD_T_ from PD_AR_ with a high degree of accuracy. Again, this result points to differential neural mechanisms in respective PD subtypes. It should be noted that the accuracy of a classifier is not a standardized estimate for its effect size. Classification accuracies can be lower for a single ON FC decoding, but may increase when data is summarized, i.e., averaged together within each subtype. This does not necessarily translate to increased statistical power because accuracy estimates are based on fewer data points, which usually increases variability. Future studies should; therefore, investigate the possibility of combining multivariate and univariate techniques to obtain new information about brain mechanisms affecting PD subtypes [64].

### Study Limitations and future directions

Cross-sectional and longitudinal studies, aimed at validating the integrity of ON FC, are needed to provide deeper insights into the relationships between olfactory and motor impairments in PD. Extensive and complex neurodegenerative processes may take place across multiple brain regions in PD subjects with mild cognitive impairments (MCI). Illuminating the characteristics of olfactory dysfunction in patients with PD-MCI will be critical in expanding our knowledge of PD pathophysiology. Specifically, it may help us prepare early intervention strategies for mitigating PD dementia development. The framework of this paper could be further expanded to better characterize PD subtypes in terms of a unified FC metric. For example, by combining intrinsic FC (identified using rs-fMRI) with task dependent FC—identified using odor-thresholds, odor-discrimination, and/or odor-identification in task fMRI paradigms.

## CONCLUSION

Alterations in PD olfactory function appears to produce distinct ON FC patterns compared to healthy controls. ON FC impairment is relatively higher in the PD_AR_ subtype, compared to the ON FC in CN and PD_T_ patients (after adjusting for GM volumes and cognitive performance). Note that the absence of significant GM volume differences in the ON should not be interpreted as PD having no effects on GM volumes. Our data clearly establishes a correlation between olfactory and motor symptoms in PD, and suggests that PD-related olfactory dysfunction has the potential to serve as a novel biomarker for a precursory assessment of PD severity. Previously, using the same study cohort, we clarified the profile of the DMN in PD_AR_ and PD_T_ [30]. Impairments in the posterior parts of the DMN were hypothesized to increase the risks of developing cognitive deficits in PD_AR_. Similarly, ON FC is impaired to a larger extent in PD_AR_, thus raising the possibility that olfactory impairment may be used as a predictive biomarker for cognitive decline and neurodegeneration [65]. Nevertheless, future studies aimed at validating these findings (as well as longitudinal investigations into the functional integrity of the ON FC in cognitively normal PD patients) may provide greater insight into the mechanisms underlying the relationships between olfaction, motor symptoms and cognitive decline in PD subtypes [66].

## ACKNOWLEDGEMENTS

This work was supported by the NINDS grant NS060722, National Institute of Aging grant AG027771, the HMC GCRC (NIH M01RR10732), GCRC Construction Grants (C06RR016499), the Leader Family Foundation, and the Department of Radiology, Penn State College of Medicine. This project also received partial funding from a grant with the Pennsylvania Department of Health using Tobacco CURE Funds.

## REFERENCES

1. Martí, M.J. and E. Tolosa, New guidelines for diagnosis of Parkinson disease. Nature Reviews Neurology, 2013. 9(4): p. 190–191.

2. Lees, A.J., J. Hardy, and T. Revesz, Parkinson’s disease. Lancet, 2009. 373(9680): p. 2055–66.

3. Zetusky, W.J., J. Jankovic, and F.J. Pirozzolo, The heterogeneity of Parkinson’s disease: clinical and prognostic implications. Neurology, 1985. 35(4): p. 522–6.

4. Jankovic, J., et al., Variable expression of Parkinson’s disease: a base-line analysis of the DATATOP cohort. The Parkinson Study Group. Neurology, 1990. 40(10): p. 1529–34.

5. Lewis, S., et al., Heterogeneity of Parkinson’s disease in the early clinical stages using a data driven approach. Journal of Neurology, Neurosurgery & Psychiatry, 2005. 76(3): p. 343–348.

6. Eggers, C., et al., Parkinson subtypes progress differently in clinical course and imaging pattern. PloS one, 2012. 7(10): p. e46813.

7. Helmich, R.C., et al., Cerebral causes and consequences of parkinsonian resting tremor: a tale of two circuits? Brain: a journal of neurology, 2012. 135(Pt 11): p. 3206–26.

8. Lewis, M.M., et al., Differential involvement of striato-and cerebello-thalamo-cortical pathways in tremor-and akinetic/rigid-predominant Parkinson’s disease. Neuroscience, 2011. 177: p. 230–9.

9. Zaidel, A., et al., Akineto-rigid vs. tremor syndromes in Parkinsonism. Current opinion in neurology, 2009. 22(4): p. 387–93.

10. Burn, D.J., Ten steps to identify atypical parkinsonism. Journal of neurology, neurosurgery, and psychiatry, 2006. 77(12): p. 1299.

11. Burn, D.J., et al., Motor subtype and cognitive decline in Parkinson’s disease, Parkinson’s disease with dementia, and dementia with Lewy bodies. Journal of neurology, neurosurgery, and psychiatry, 2006. 77(5): p. 585–9.

12. Schapira, A.H., K.R. Chaudhuri, and P. Jenner, Non-motor features of Parkinson disease. Nature Reviews Neuroscience, 2017. 18(7): p. 435.

13. Doty, R.L., Olfactory dysfunction in Parkinson disease. Nat Rev Neurol, 2012. 8(6): p. 329–39.

14. Doty, R.L., Olfaction in Parkinson’s disease and related disorders. Neurobiol Dis, 2012. 46(3): p. 527–52.

15. Kang, S.H., et al., The combined effect of REM sleep behavior disorder and hyposmia on cognition and motor phenotype in Parkinson’s disease. Journal of the neurological sciences, 2016. 368: p. 374–378.

16. Siderowf, A., et al., Impaired olfaction and other prodromal features in the Parkinson At-Risk Syndrome Study. Movement Disorders, 2012. 27(3): p. 406–412.

17. Postuma, R. and J.-F. Gagnon, Cognition and olfaction in Parkinson’s disease. Brain: a journal of neurology, 2010. 133(12): p. e160–e160.

18. Fullard, M.E., et al., Olfactory impairment predicts cognitive decline in early Parkinson’s disease. Parkinsonism & related disorders, 2016. 25: p. 45–51.

19. Hoyles, K. and J.C. Sharma, Olfactory loss as a supporting feature in the diagnosis of Parkinson’s disease: a pragmatic approach. Journal of neurology, 2013. 260(12): p. 2951–2958.

20. Berg, D., et al., MDS research criteria for prodromal Parkinson’s disease. Movement Disorders, 2015. 30(12): p. 1600–1611.

21. Heinzel, S., et al., Update of the MDS research criteria for prodromal Parkinson’s disease. Movement Disorders, 2019. 34(10): p. 1464–1470.

22. Helmich, R.C., et al., Pallidal dysfunction drives a cerebellothalamic circuit into Parkinson tremor. Annals of neurology, 2011. 69(2): p. 269–281.

23. Dauer, W. and S. Przedborski, Parkinson’s disease: mechanisms and models. Neuron, 2003. 39(6): p. 889–909.

24. Ponsen, M.M., et al., Idiopathic hyposmia as a preclinical sign of Parkinson’s disease. Annals of Neurology: Official Journal of the American Neurological Association and the Child Neurology Society, 2004. 56(2): p. 173–181.

25. Baba, T., et al., Association of olfactory dysfunction and brain. Metabolism in Parkinson’s disease. Movement Disorders, 2011. 26(4): p. 621–628.

26. Tissingh, G., et al., Loss of olfaction in de novo and treated Parkinson’s disease: possible implications for early diagnosis. Movement disorders: official journal of the Movement Disorder Society, 2001. 16(1): p. 41–46.

27. Doty, R.L., et al., Bilateral olfactory dysfunction in early stage treated and untreated idiopathic Parkinson’s disease. Journal of Neurology, Neurosurgery & Psychiatry, 1992. 55(2): p. 138–142.

28. Poewe, W., et al., Parkinson disease. Nature reviews Disease primers, 2017. 3(1): p. 1–21.

29. Hummel, T., et al., Olfactory FMRI in patients with Parkinson’s disease. Frontiers in Integrative Neuroscience, 2010. 4: p. 125.

30. Karunanayaka, P.R., et al., Default mode network differences between rigidity-and tremor-predominant Parkinson’s disease. Cortex; a journal devoted to the study of the nervous system and behavior, 2016. 81: p. 239–50.

31. Goodyear, M.D., K. Krleza-Jeric, and T. Lemmens, The declaration of Helsinki, 2007, British Medical Journal Publishing Group.

32. Calne, D., B. Snow, and C. Lee, Criteria for diagnosing Parkinson’s disease. Annals of Neurology: Official Journal of the American Neurological Association and the Child Neurology Society, 1992. 32(S1): p. S125–S127.

33. Schiess, M.C., et al., Parkinson’s disease subtypes: clinical classification and ventricular cerebrospinal fluid analysis. Parkinsonism & related disorders, 2000. 6(2): p. 69–76.

34. Seubert, J., et al., Statistical localization of human olfactory cortex. NeuroImage, 2013. 66: p. 333–342.

35. Tobia, M.J., Q.X. Yang, and P. Karunanayaka, Intrinsic intranasal chemosensory brain networks shown by resting-state functional MRI. Neuroreport, 2016. 27(7): p. 527–31.

36. Karunanayaka, P., M.J. Tobia, and Q.X. Yang, Age-related resting-state functional connectivity in the olfactory and trigeminal networks. Neuroreport, 2017. 28(15): p. 943–948.

37. Lu, J., et al., Functional Connectivity between the Resting-State Olfactory Network and the Hippocampus in Alzheimer’s Disease. Brain sciences, 2019. 9(12).

38. Brodersen, K.H., et al., Generative embedding for model-based classification of fMRI data. PLoS Comput Biol, 2011. 7(6): p. e1002079.

39. Fishman, P.S., Paradoxical aspects of parkinsonian tremor. Movement Disorders, 2008. 23(2): p. 168–173.

40. Witt, M., et al., Biopsies of olfactory epithelium in patients with Parkinson’s disease. Movement disorders: official journal of the Movement Disorder Society, 2009. 24(6): p. 906–914.

41. Haehner, A., et al., Substantia nigra fractional anisotropy changes confirm the PD at-risk status of patients with idiopathic smell loss. Parkinsonism & related disorders, 2018. 50: p. 113–116.

42. Domellöf, M.E., et al., Olfactory dysfunction and dementia in newly diagnosed patients with Parkinson’s disease. Parkinsonism & related disorders, 2017. 38: p. 41–47.

43. Marin, C., et al., Olfactory dysfunction in neurodegenerative diseases. Current allergy and asthma reports, 2018. 18(8): p. 42.

44. White, T.L., A.F. Sadikot, and J. Djordjevic, Metacognitive knowledge of olfactory dysfunction in Parkinson’s disease. Brain and cognition, 2016. 104: p. 1–6.

45. Dirkx, M.F., et al., The cerebral network of Parkinson’s tremor: an effective connectivity fMRI study. Journal of Neuroscience, 2016. 36(19): p. 5362–5372.

46. Mundinano, I.-C., et al., Increased dopaminergic cells and protein aggregates in the olfactory bulb of patients with neurodegenerative disorders. Acta Neuropathol, 2011. 122(1): p. 61.

47. Cole, M.W., et al., Intrinsic and task-evoked network architectures of the human brain. Neuron, 2014. 83(1): p. 238–251.

48. Ito, T., et al., Discovering the computational relevance of brain network organization. Trends in cognitive sciences, 2020. 24(1): p. 25–38.

49. Braak, H. and E. Braak, Evolution of neuronal changes in the course of Alzheimer’s disease. J Neural Transm Suppl, 1998. 53: p. 127–40.

50. Braak, H., et al., Cognitive status correlates with neuropathologic stage in Parkinson disease. Neurology, 2005. 64(8): p. 1404–10.

51. Wang, J., et al., Association of olfactory bulb volume and olfactory sulcus depth with olfactory function in patients with Parkinson disease. American journal of neuroradiology, 2011. 32(4): p. 677–681.

52. Deco, G., V.K. Jirsa, and A.R. McIntosh, Emerging concepts for the dynamical organization of resting-state activity in the brain. Nature Reviews Neuroscience, 2011. 12(1): p. 43–56.

53. Adachi, Y., et al., Functional connectivity between anatomically unconnected areas is shaped by collective network-level effects in the macaque cortex. Cerebral cortex, 2012. 22(7): p. 1586–1592.

54. Koch, M.A., D.G. Norris, and M. Hund-Georgiadis, An investigation of functional and anatomical connectivity using magnetic resonance imaging. NeuroImage, 2002. 16(1): p. 241–250.

55. Honey, C.J., et al., Network structure of cerebral cortex shapes functional connectivity on multiple time scales. Proceedings of the National Academy of Sciences, 2007. 104(24): p. 10240–10245.

56. Westermann, B., et al., Functional imaging of the cerebral olfactory system in patients with Parkinson’s disease. Journal of Neurology, Neurosurgery & Psychiatry, 2008. 79(1): p. 19–24.

57. Parkkinen, L., et al., Can olfactory bulb biopsy be justified for the diagnosis of Parkinson’s disease? Comments on” olfactory bulb [alpha]-synucleinopathy has high specificity and sensitivity for Lewy body disorders”. Acta Neuropathol, 2009. 117(2): p. 213.

58. Brodoehl, S., et al., Decreased olfactory bulb volume in idiopathic Parkinson’s disease detected by 3.0-Tesla magnetic resonance imaging. Movement Disorders, 2012. 27(8): p. 1019–1025.

59. Mueller, A., et al., Olfactory bulb volumes in patients with idiopathic Parkinson’s disease a pilot study. Journal of neural transmission, 2005. 112(10): p. 1363–1370.

60. Wattendorf, E., et al., Olfactory impairment predicts brain atrophy in Parkinson’s disease. Journal of Neuroscience, 2009. 29(49): p. 15410–15413.

61. Wu, X., et al., Correlation between progressive changes in piriform cortex and olfactory performance in early Parkinson’s disease. European Neurology, 2011. 66(2): p. 98–105.

62. Lee, J.E., et al., Olfactory performance acts as a cognitive reserve in non-demented patients with Parkinson’s disease. Parkinsonism & related disorders, 2014. 20(2): p. 186–191.

63. Tessitore, A., et al., Resting-state brain connectivity in patients with Parkinson’s disease and freezing of gait. Parkinsonism & related disorders, 2012. 18(6): p. 781–7.

64. Weichwald, S., et al., Causal interpretation rules for encoding and decoding models in neuroimaging. NeuroImage, 2015. 110: p. 48–59.

65. Kawasaki, I., et al., Loss of awareness of hyposmia is associated with mild cognitive impairment in Parkinson’s disease. Parkinsonism & related disorders, 2016. 22: p. 74–79.

66. Aarsland, D., et al., Cognitive decline in Parkinson disease. Nature Reviews Neurology, 2017. 13(4): p. 217–231.

