## Supplementary Table S1 for "IMPAIRED OLFACTORY NETWORK FUNCTIONAL CONNECTIVITY IN PARKINSON’S DISEASE: A NOVEL MARKER FOR DISEASE PROGRESSION"

### (SUPPLEMENTARY MATERIAL)

**Abbreviated title:** Impaired ON FC in PD

Prasanna Karunanayaka[1], Jiaming Lu[1,2], Mechelle M. Lewis[3], Rommy Elyan[1], Qing X. Yang[1,4], Paul J. Eslinger[1,4], Xuemei Huang[3]

[1]Department of Radiology, The Pennsylvania State University College of Medicine, Hershey, PA, USA

[2]Drum Tower Hospital, Medical School of Nanjing University, Nanjing, China

[3]Department of Neurology, The Pennsylvania State University College of Medicine, Hershey, PA, USA

[4]Department of Neurosurgery, The Pennsylvania State University College of Medicine, Hershey, PA, USA

| PD subtypes | Total tremor scores | Mean Tremor score | Total AR scores | Mean AR score | Mean Tremor/AR ratio |
|---|---|---|---|---|---|
| **PD$_{AR}$ patients** | | | | | |
| 4 | 0 | 0.00 | 10 | 0.71 | 0.00 |
| 7 | 4 | 0.44 | 11 | 0.79 | 0.57 |
| 16 | 1 | 0.11 | 8 | 0.57 | 0.19 |
| 22 | 0 | 0.00 | 15 | 1.07 | 0.00 |
| 30 | 1 | 0.11 | 11 | 0.79 | 0.14 |
| 36 | 0 | 0.00 | 30 | 2.14 | 0.00 |
| 45 | 10 | 1.11 | 33 | 2.36 | 0.47 |
| 59 | 6 | 0.67 | 11 | 0.79 | 0.85 |
| 67 | 4 | 0.44 | 18 | 1.29 | 0.35 |
| 68 | 0 | 0.00 | 16 | 1.14 | 0.00 |
| 73 | 0 | 0.00 | 6 | 0.43 | 0.00 |
| 92 | 5 | 0.56 | 17 | 1.21 | 0.46 |
| 104 | 5 | 0.56 | 26 | 1.86 | 0.30 |
| 129 | 3 | 0.33 | 23 | 1.64 | 0.20 |
| 183 | 0 | 0.00 | 10 | 0.71 | 0.00 |
| 187 | 0 | 0.00 | 6 | 0.43 | 0.00 |
| 190 | 0 | 0.00 | 30 | 2.14 | 0.00 |
| **PD$_{T}$ patients** | | | | | |
| 23 | 6 | 0.67 | 6 | 0.43 | 1.56 |
| 31 | 7 | 0.78 | 9 | 0.64 | 1.21 |
| 40 | 3 | 0.33 | 3 | 0.21 | 1.56 |
| 54 | 6 | 0.67 | 5 | 0.36 | 1.87 |
| 74 | 11 | 1.22 | 12 | 0.86 | 1.43 |
| 81 | 3 | 0.33 | 4 | 0.29 | 1.17 |
| 85 | 3 | 0.33 | 5 | 0.36 | 0.93 |
| 87 | 5 | 0.56 | 1 | 0.07 | 7.78 |
| 94 | 7 | 0.78 | 3 | 0.21 | 3.63 |

| | | | | | |
|---|---|---|---|---|---|
| 100 | 11 | 1.22 | 7 | 0.50 | 2.44 |
| 105 | 4 | 0.44 | 5 | 0.36 | 1.24 |
| 111 | 4 | 0.44 | 6 | 0.43 | 1.04 |
| 178 | 7 | 0.78 | 6 | 0.43 | 1.81 |
| 181 | 9 | 1.00 | 15 | 1.07 | 0.93 |
| 182 | 5 | 0.56 | 4 | 0.29 | 1.94 |

**Table S1**. Full summary of UPDRS scores for $PD_{AR}$ and $PD_T$. AR indicates akinetic-rigidity subtype. T indicates tremor-predominant subtype.
